# A distributed theta network of error generation and processing in aging

**DOI:** 10.1101/2023.06.28.546844

**Authors:** Vasil Kolev, Michael Falkenstein, Juliana Yordanova

**Author notes:** **Corresponding author:** Prof. Dr. Juliana Yordanova, Institute of Neurobiology, Bulgarian Academy of Sciences, Acad. G. Bonchev str., bl. 23, 1113 Sofia, Bulgaria.

## Abstract

Based on previous concepts that a distributed theta network with a central “hub” in the medial frontal cortex is critically involved in movement regulation, monitoring, and control, the present study explored the involvement of this network in error processing with advancing age in humans. For that aim, the oscillatory neurodynamics of motor theta oscillations was analyzed at multiple cortical regions during correct and error responses in a sample of older adults.

Response-related potentials (RRPs) of correct and incorrect reactions were recorded in a four-choice reaction task. RRPs were decomposed in the time-frequency domain to extract oscillatory theta activity. Motor theta oscillations at extended motor regions were analyzed with respect to power, temporal synchronization, and functional connectivity.

Major results demonstrated that errors had pronounced effects on motor theta oscillations at cortical regions beyond the medial frontal cortex by being associated with (1) theta power increase in the hemisphere contra-lateral to the movement, (2) suppressed spatial and temporal synchronization at pre-motor areas contra-lateral to the responding hand, (2) inhibited connections between the medial frontal cortex and sensorimotor areas, and (3) suppressed connectivity and temporal phase-synchronization of motor theta networks in the posterior left hemisphere, irrespective of the hand, left, or right, with which the error was made.

These findings reveal distributed effects of errors on motor theta oscillations in older subjects and support the hypothesis that error processing operates on a network level. They confirm the presence of aging-dependent functional disengagement of the medial frontal region and suggest that difficulties in controlling the focus of motor attention and response selection contribute to performance impairment in old individuals.

## 1. Introduction

A negative brain potential with midline fronto-central maximum, called error negativity (Ne; Falkenstein et al. 1990, 1991) or error-related negativity (ERN; Gehring et al. 1993) is an established neurophysiological marker of error processing in humans (rev. Vidal et al. 2022; Fu et al. 2023). The Ne is specifically elicited or substantially enhanced by performance errors and is generated by medial frontal structures (rev. Vidal et al. 2022). Although different models have been proposed for Ne in relation to detecting a mismatch between the planned and the actual action (Falkenstein et al. 2001; Dehaene 2018), processing of conflict between concurrent response options (Carter et al. 1998; Botvinick et al. 2001, 2004), or detecting unpredicted behavioural outcomes (Holroyd and Coles 2002; Alexander and Brown 2011; Silvetti et al. 2011), all models support the role of Ne for behavioural monitoring, control and adaptation (Carter et al. 1998; Ullsperger and von Cramon 2001, 2004; Vidal et al. 2022).

The Ne has been intensively studied to explore the neurophysiologic mechanisms of error processing in aged population. It has been consistently reported that the Ne is significantly reduced in older subjects (Band and Kok 2000; Falkenstein et al. 2001; Mathalon et al. 2003; Nieuwenhuis et al. 2002; Kolev et al. 2005) suggesting an aging-related deficiency in performance monitoring. To understand more precisely the origin of Ne reduction with aging, time-frequency decomposition methods have been applied. These approaches have revealed that the Ne reduction with aging is associated with a substantial decrease in the power and response-locked synchronization of medial frontal theta oscillations (Kolev et al. 2009).

Interestingly, despite the aging-related Ne suppression, no differences in error rate, correction rate, or post-error slowing have been found between young and older adults in choice-reaction sensorimotor tasks (Falkenstein et al. 2001; Kolev et al. 2005). Similar dissociations between ongoing, adaptive, and corrective performance and age-dependent Ne alterations have been detected in other task conditions (Band and Kok 2000; Mathalon et al. 2003). Also, in contrast to young adults manifesting a strong correlation between response speed and medial frontal theta power, no such correlations have been observed in older adults (Yordanova et al. 2020). Since the age-dependent reduction of error-related signaling from the fronto-medial area appears not to be not uniquely associated with performance quality, it can be hypothesized that a network reorganization that takes place with aging (Cabeza 2002; Cabeza et al. 2002; Reuter-Lorenz 2002; Li et al. 2001; Koen and Rugg 2019) leads to different neurophysiological support of error processing involving distributed brain regions in older subjects.

This network hypothesis is substantiated by previous studies revealing that delta/theta oscillations observed during Ne are generated not only at the medial frontal area but also at the hemisphere contralateral to the responding hand during response generation (Yordanova et al. 2004, 2023). Also, analyses of response-related potentials (RRPs) during correct response production have found that RRPs at extended cortical areas contain delta/theta components in both young and older adults, similar to Ne at the medial frontal cortex (Popovych et al. 2016; Liu et al. 2017; Yordanova et al. 2020). Moreover, the activity of the medial frontal cortex during all phases of movement production (preparation, initiation, and execution) emerges in close association with motor cortical oscillations (Urbano et al. 1996, 1998a, b). This co-activation exists also for error responses and is mainly supported by alternating oscillatory patterns from the theta frequency band (Luu and Tucker 2001; Yordanova et al. 2004, 2023). These observations support the notion that a distributed theta system in the brain with a central coordinating “hub” in the medial frontal regions (Cohen 2011, 2014) orchestrates cyclically brain computations in distant regions in relation to motor response control and execution (Duprez et al. 2020). Thus, if distributed theta networks are critically involved in movement regulation, monitoring, and control, it can be further hypothesized that altered error processing in old subjects may be reflected by error-related theta activity at multiple regions. To test this hypothesis, the present study explored the oscillatory neurodynamics of motor theta oscillations over extended cortical areas during correct and error movement generation in a group of older people.

## 2. Methods

### 2.1. Subjects

A total of 32 subjects were included in the study (16 young and 16 older subjects). They were selected from a larger sample also used in a study of correct motor potentials (Yordanova et al. 2020). Due to the application of an adopted stringent criterion for error analysis, the final samples accepted for analysis included 10 young adults (5f, mean age 22.5 years, SE ±1.5) and 11 older adults (6 females, mean age 58.3 years, SE = ±2.1). All participants were healthy, without a history of neurologic, psychiatric, chronic somatic, or hearing problems, with normal or corrected-to-normal vision, and were involved in social and working activities. They were right-handed and took no medication during the experimental sessions. Experiments were approved by the local ethic committee of the Leibniz Institute for Working Environment and Human Factors, Dortmund, Germany. Prior to engaging with the study, all participants gave informed consent in line with the Declaration of Helsinki. All older adults were in an active working stage of life. As detailed below, in the present study, a general analysis of data will conducted using the two age groups, and complete reports will include only the group of older adults.

### 2.2. Task

A four-choice reaction task (CRT) was employed as described in Yordanova et al. (2020). Four stimulus types represented by the letters A, E, I, and O were delivered randomly with an equal probability in separate experimental blocks and had to be responded to with the left middle, left index, right index, and right middle fingers, respectively. A total of 200 stimuli were presented in each block. Response force was measured by sensometric tensors while subjects produced a flexion with each of the four fingers. The CRT was performed in two modalities - auditory and visual. Auditory stimuli (duration 300 ms, intensity 67 dB SPL) were delivered via headphones binaurally. Visual stimuli (duration 300 ms) were shown in the middle of a monitor placed 1.5 m in front of the subject. Inter-stimulus intervals varied randomly between 1440 and 2160 ms (mean 1800 ms). In the case of slow responses, a feedback tone was delivered at 700 ms after stimulus onset and had to be avoided by speeding up reactions. A total of nine auditory and nine visual CRT blocks were performed by each participant. Experimental blocks were randomized and sequences of auditory and visual blocks were counterbalanced across participants.

### 2.3. Data recording and processing

Data recording and analysis followed the procedures described in Yordanova et al. (2020). Data from all nine sessions in each modality were used. EEG was recorded from 64 channels with Cz as reference, with frequency limits of 0.1–70 Hz, and a sampling rate of 250 Hz. EEG traces were visually inspected for gross electrooculogram (EOG) and electromyogram (EMG) artifacts. Contaminated trials were discarded along with EEG traces exceeding ±100 µV. The accepted trials were corrected using a linear regression method (Gratton et al. 1983). Mechanograms from each finger were recorded for analysis of response correctness and speed. Data processing was performed using Brain Vision Analyzer 2.2.2 (Brain Products GmbH, Gilching, Germany).

### 2.4. Response-related potentials

Response-related potentials (RRPs) were computed with a trigger corresponding to a threshold level of 5 N in the mechanogram, thus discarding incomplete responses. For each stimulus-response type (SR1, SR2, SR3, and SR4), between 20 and 30 artifact-free error trials from each individual were used. A randomized inclusion procedure was used to equalize the number of error and correct trials for each subject and SR type. Responses with left-hand and right-hand fingers were combined to produce RRPs for left- and right-hand responses separately. Thus, between 40 and 60 single sweeps for each participant in each modality (auditory and visual) for each hand (left and right) were used for RRP analysis of correct and incorrect responses. All RRP analyses were performed after current source density (CSD) transform of the signals (e.g., Babiloni et al. 1996; Nunez et al. 1997; Perrin et al. 1989) providing for a reference-free evaluation. The exact mathematical procedure is presented in detail in Perrin et al. (1989).

### 2.5. Time–frequency decomposition

Time-frequency (TF) analysis of RRPs was performed by means of a continuous wavelet transform (CWT). Details of the mathematical procedure are presented in Yordanova et al. (2020). The analysis was performed in the frequency range of 0.1-16 Hz with a central frequency at 0.4 Hz intervals.

To achieve a reliable analysis of low-frequency components in the time-frequency domain and avoid possible edge effects, 4096 ms-long epochs were used, with the moment of response execution (5 N) being in the center of the analysis epoch. A baseline of 600-800 ms before the response was used. TF decomposition was performed on CSD-transformed single-sweep RRPs. Basing on previous observations of TF plots of RRPs, the theta layer was extracted with a central frequency of 5.5 Hz.

#### 2.5.1. Total power

Total power (TOTP) comprises the phase-locked and non-phase-locked fractions of the signal. It was measured to represent the total energy of response-related oscillations. For each trial, the time-varying power in the theta band was calculated by squaring the absolute value of the convolution of the signal with the complex wavelet.

#### 2.5.2. Temporal synchronization

The phase synchronization across trials was measured by means of the phase-locking factor (PLF, e.g., Lachaux et al. 1999; Tallon-Baudry et al. 1997). The PLF provides a measure of synchronization of oscillatory activity independently of the signal’s amplitude. The values of PLF yield a number between 0 and 1 determining the degree of between-sweep phase-locking, where 1 indicates perfect phase alignment across trials and values close to 0 reflect the highest phase variability.

#### 2.5.3. Spatial synchronization

To assess spatial synchronization across distant cortical regions, the phase-locking value (PLV) was used (Cohen 2015). PLV measures the extent to which oscillation phase angle differences between electrodes are consistent over trials at each time/frequency point. PLVs were computed for the theta TF scale at each time point and trial (computation details in Yordanova et al. 2017, 2023). For PLV analysis 35 electrodes were used (F3, Fz, F4, FC5, FC3, FC1, FCz, FC2, FC4, FC6, T7, C5, C3, C1, Cz, C2, C4, C6, T8, CP5, CP3, CP1, CPz, CP2, CP4, CP6, P3, Pz, P4, PO5, POz, PO6, O1, Oz, and O2). PLV was computed for each pair of electrodes, resulting in a total of 595 pairs for each subject, modality, hand (left and right), and response type (correct and error).

### 2.6. Parameters

The following TF parameters were computed: TOTP and PLF at 64 electrodes, and PLV of 595 electrode pairs. In addition, two other parameters were introduced based on the pair-wise PLV measures. To identify regions with maximal connectedness with all other cortical regions (“hubs”) during response production, the mean of all pairs (n = 34) was computed for every single electrode (regional PLV, R-PLV). Also, a separate analysis used PLV measures of pairs guided by the medial fronto-central electrode FCz (FCz-PLV). This measure aimed at specifically assessing the connectivity of the acknowledged response monitoring medial frontal region during correct and error response generation.

For all TF parameters (TOTP, PLF, R-PLV, and FCz-PLV) the maximal value was identified in the latency range of -300 to +300 ms around the moment of response execution. The parameter was measured as the mean magnitude value within -24 to +24 ms around the maximum. In addition, the peak latency of the maximum was measured according to the moment of response production. For statistical evaluation, measures of TOTP were log10-transformed.

### 2.7. Statistical analyses

Response-related TF parameters were analyzed at motor cortical areas contra-lateral and ipsi-lateral to the responding hand. Three regions of interest were used – fronto-central, central and centro-parietal. A repeated-measures ANOVA design was applied with within-subjects variables Accuracy (Correct vs. Error) and Modality (Auditory vs. Visual). Additional within-subjects factors were included to analyze topographic effects – Region (fronto-central FC3/FCz/FC4 vs. central C3/Cz/C4 vs. centro-parietal CP3/CPz/CP4) and Laterality (left hemisphere FC3/C3/CP3 vs. midline FCz/Cz/CPz vs. right hemisphere FC4/C4/CP4). Only for FCz-PLV, did the Laterality variable include two levels – left hemisphere and right hemisphere electrodes. Analyses were performed separately for the left- and right-hand responses to control for the effects of hemispheres contra- and ipsi-lateral to the response that were opposite for the left- and right-hand responses. Correct and incorrect reaction times (RT) of right- and left-hand responses as well as error rates were analyzed in an Accuracy x Modality x Response Side (Left hand vs. Right hand) ANOVA design. Greenhouse-Geisser corrected p-values are reported. Pearson correlations were computed to study relevant relationships. In the general analysis, Age was included in all ANOVAs as a between-subjects factor.

## 3. Results

### General Analysis

For TOTP, PLF and R-PLV theta parameters, significant Age x Accuracy x Topography interactions were yielded in both the left- and right-hand analyses: Age x Accuracy x Region: F(2/38) = 3.6 – 6.9, p = 0.05 – 0.004; Age x Accuracy x Laterality (F(2/38) = 5.3 – 13.3, p = 0.01 – 0.001); Age x Accuracy x Region x Laterality (F(4/76) = 3.5 – 7.6, p = 0.02 – 0.001). For FCz-PLV these interactions were significant only for left-hand responses: Age x Accuracy x Region: F(2/38) = 4.6, p = 0.03; Age x Accuracy x Laterality (F(1/19) = 5.4, p = 0.03); Age x Accuracy x Region x Laterality (F(2/38) = 5.3, p = 0.01). These complex interactions reflected a substantial difference that characterized the involvement of motor theta oscillations in the two age groups. For that reason, the detailed exploration of error-related effects in young adults is presented in a separate study (Yordanova et al., 2023). RT was significantly longer in older than young adults (F(1/19), which was valid for responses with each hand – Fig. 1. In the two age groups, the slower left-hand correct responses were associated with faster errors, whereas the faster right-hand correct responses were associated with slower errors (Accuracy x Side, F(1/19) = 20.9, p < 0.001; Accuracy x Side x Age, F(1/19) = 1.4, p > 0.2). Error rate did not depend on Age (F1/19) = 0.19, p > 0.6), and was higher for the left hand in two groups (Side, F(1/19) = 9.9, p = 0.005; Age x Side, F(1/19) = 0.8, p > 0.4). In the following, the results of only the older adults are described, and the results of young subjects are only illustrated for comparative purposes (Fig. 1, 2 – right panels) and additionally briefly outlined in the Supplementary Information.

**Figure 1.**
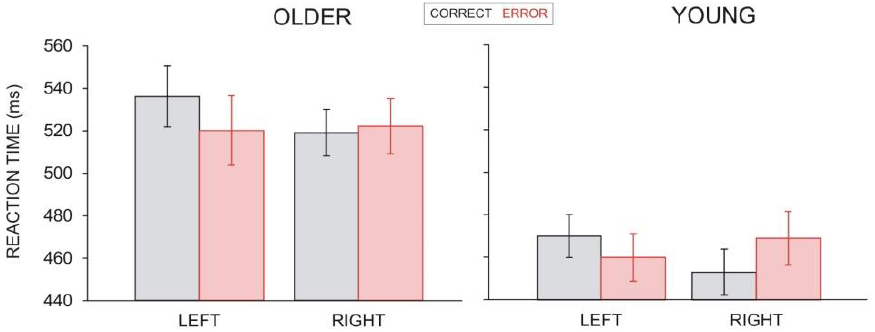
Group mean reaction times for correct and error responses produced with the left and the right hand. Left panel – OLDER adults. Right panel – YOUNG adults (with modifications from Yordanova et al., 2023).

**Figure 2.**
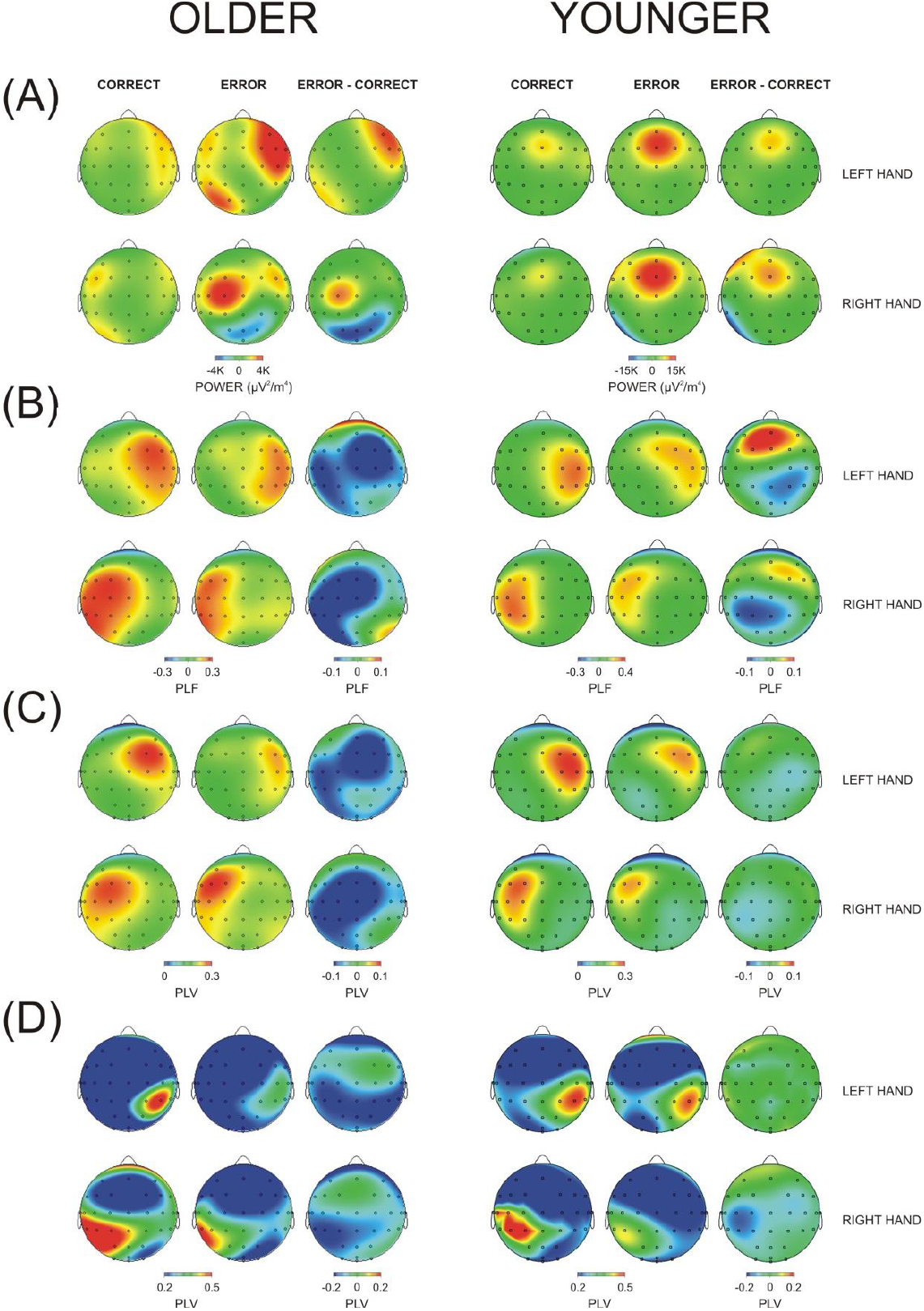
Topography maps for correct, error and error minus correct difference at the time of maximal expression of the theta TF component (3.5-7 Hz) of response-related potentials elicited by left- and right-hand motor responses (A) Total power TOTP, (B) Temporal synchronization (PLF), (C) Region-specific connectedness R-PLV, (D) FCz-guided synchronization FCz-PLV. Left panel – OLDER adults. Right panel – YOUNG adults (with modifications from Yordanova et al., 2023). Response onset at 0 ms.

### Analysis of Older Adults

#### 3.1. Performance

RT did not depend on Modality, Accuracy, and Side (F(1/10) < 1.2, p > 0.2). However, there was a significant Accuracy x Side interaction (F(1/10) = 5.3, p = 0.04) – Fig. 1(left). Consistent with the right-handedness of the subjects, correct RTs were faster for right-(mean 517 ms) than left-hand (mean 537 ms) responses (Side, F(1/10) = 11.5, p = 0.005). For the faster right hand, errors appeared slower than correct responses (522 vs. 517 ms, ns.), whereas, for the slower left hand, errors were faster than correct responses (537 vs. 520 ms, p = 0.04). Error rate was higher for the left hand (Side, F(1/10) = 8.8, p = 0.01).

#### 3.2. Response-related theta oscillations

##### 3.2.1. Theta TOTP

Figure 2A - left panel demonstrates that the power of motor-related theta oscillations was distributed asymmetrically to the hemisphere contra-lateral to the response, which was significant for the left hand (Laterality, F(2/20) = 9.8/3.3, p = 0.007/0.07 for left- and right-hand responses, respectively). Errors were associated with an overall increase in theta TOTP (Accuracy, F(1/10) = 12.1/8.5, p = 0.006/0.015). This increase was pronounced at the right pre-motor areas for the left-hand errors (Accuracy x Laterality, F(2/20) = 13.6, p < 0.001) and at the left motor area for right-hand errors (Fig. 2A - left panel), but this latter effect was not significant.

##### 3.2.2. Theta PLF

As illustrated in Fig. 2B – left panel, response-related theta PLF was significantly stronger at contra- than ipsilateral regions (Laterality, F(2/20) = 29.8/4.4, p = 0.001/0.03 for left-and right-hand responses, respectively). The difference between contra- and ipsilateral hemispheres was highly significant for both hands (F(1/10) = 56.5/23.9, p < 0.0001).

As verified by significant main and interactive effects (Accuracy, F(1/10) = 38.4/17.1, p < 0.001/0.002; Accuracy x Laterality, F(2/20) = 11.8/13.7, p < 0.001; Accuracy x Region x Laterality, F(4/40) = 3.0/3.6, p = 0.05/0.01), errors were associated with a significant decrease of theta PLF over the contra-lateral and midline fronto-central and central regions – Fig. 2B - left panel. Notably, for both response sides, this contra-lateral theta PLF suppression was accompanied by a significant decrease of the temporal synchronization over the left hemisphere. Accordingly, for left-hand responses, the simple Accuracy effect was significant at FC4, FCz, C4, Cz, C3 and CP3 electrodes (F(1/10) = 8.3 – 31.8, p = 0.02 – 0.001). For right-hand responses, the simple Accuracy effect was significant at FC3, FCz, C3, Cz, and CP3 electrodes (F(1/10) = 7.5 – 20.1, p = 0.02 – 0.001).

##### 3.2.3. Theta R-PLV

Figure 2C - left panel demonstrates that during both left- and right-hand responses, the region-specific connectivity with all other cortical areas was significantly stronger for the hemisphere contra-than ipsilateral to the response (Laterality, F(2/20) = 19.4/12.4, p < 0.001).

Similar to theta PLF, theta R-PLV was significantly reduced during both left- and right-hand errors at the contra-lateral and midline fronto-central and central regions as well as at the left posterior electrodes (Accuracy, F(1/10) = 12.3/9.02, p < 0.006/0.01; Accuracy x Laterality, F(2/20) = 6.4/4.6, p = 0.01/0.02; Accuracy x Region x Laterality, F(4/40) = 5.1/4.6, p = 0.007/0.01). For left-hand responses, the simple Accuracy effect was significant at FC4, FCz, Cz, C3 and CP3 electrodes (F(1/10) = 4.8 – 14.1, p = 0.05 – 0.004). For right-hand responses, the simple Accuracy effect was significant at FC3, FCz, C3, Cz, and CP3 electrodes (F(1/10) = 5.7 – 10.9, p = 0.04 – 0.008).

##### 3.2.4. Theta FCz-PLV

Figure 2D - left panel demonstrates that during both left- and right-hand hand responses, theta oscillations were synchronized between FCz and centro-parietal regions (Region, F(2/20 = 5.9/3.8, p = 0.01/0.04). Also, the FCz-guided spatial synchronization was significantly stronger at the hemisphere contra-lateral than ipsilateral to the response (Laterality, F(1/10) = 16.7/5.4, p = 0.002/0.04). The contra-vs. ipsilateral difference in FCz-guided synchronization was most expressed for the central and centro-parietal regions (Region x Laterality, F(4/40) = 5.04/6.1, p = 0.04/0.01).

Errors produced a region-specific reduction of the FCz-guided synchronization. Theta FCz-PLV was suppressed by left-hand errors at the centro-parietal electrodes of both the left and the right hemisphere (Accuracy, F(1/10) = 6.6, p = 0.03; Accuracy x Region, F(2/10) = 4.5, p = 0.02; simple Accuracy effect at centro-parietal region, F(1/10) = 11.0, p = 0.008). In contrast, FCz-PLV was reduced by right-hand errors only at the contra-lateral centro-parietal region (Accuracy x Region x Laterality, F(4/40) = 6.3, p = 0.01; simple Accuracy effect at CP3, F(1/10) = 15.9, p = 0.003). These observations show that FCz-PLV is reduced by errors at contra-lateral centro-parietal regions, and additionally at the left posterior regions for left-hand errors.

#### 3.3. Correlational analyses

According to a previous study (Yordanova et al. 2020), RT of correct responses in young adults correlated negatively with theta TOTP at FCz, in contrast to old adults who did not demonstrate any relationships. We tested if theta TOTP found here to be distributed at contra-lateral regions would manifest the same associations with response speed, which would inform about preserved functional sensitivity. In older subjects, the computation of Pearson correlations revealed moderate to strong negative correlations between theta TOTP at contra-lateral electrodes and correct and error RT of the left hand (r = - 0.6/-0.8, p = 0.05 – 0.005) and the right hand (r = - 0.6/-0.7, p = 0.05 – 0.03). Another question of interest was if the error-related suppression of synchronization at different cortical regions in older adults was independent or co-existing. For PLF and R-PLV it was tested if the error-related reduction of synchronization at FCz correlated with the same effects at contra-lateral pre-motor regions. A positive correlation was found only for left-hand responses between FCz and FC4 (r = 0.63, p = 0.03).

## 4. Discussion

Based on previous concepts of a distributed theta network with a central “hub” in the medial frontal cortex (Cohen 2011, 2014) that is critically involved in movement regulation, monitoring, and control (Yordanova et al. 2004, 2020; Hoffmann et al. 2014; Duprez et al. 2020), the present study was undertaken to explore the involvement of this network in error processing in advanced age in humans. For that aim, the oscillatory neurodynamics of motor theta oscillations at multiple cortical regions was analyzed during correct and error responses in a sample of older adults.

In line with earlier studies of old subjects, an error-related suppression of temporal and spatial synchronizations of theta activity was found at the medial frontal area (Kolev et al. 2005, 2009). However, errors had pronounced effects on motor theta oscillations at cortical regions beyond the medial frontal cortex. Specifically, incorrect responses were associated with (1) theta power increase in the hemisphere contralateral to the movement, (2) suppressed spatial and temporal synchronization at contra-lateral pre-motor areas, (3) inhibited connections between the medial frontal cortex and contra-lateral sensorimotor areas, and (4) suppressed connectivity and temporal phase-synchronization of motor theta networks in the posterior left hemisphere, irrespective of the hand, left, or right, with which the error was made. Together, these findings reveal novel distributed effects of errors on motor theta oscillations in older individuals and support the hypothesis that error processing operates on a network level.

According to one major finding, theta power in older adults was overall increased after errors, particularly at contra-lateral pre-motor and motor regions. Confirming earlier studies (Kolev et al. 2005, 2009) no error-related enhancement was evident at the FCz location, in contrast with findings in young adults who manifest the most remarkable increase of error-related theta power at the frontal-central midline (Kolev et al. 2005, 2009; Yordanova et al. 2023; Fig. 2A - right panel). Since enhanced theta (4-8 Hz) oscillations centered at FCz have been associated with a variety of executive and cognitive control functions - conflict processing, detection of errors, inhibition, performance monitoring, and behavioral re-adjustment (Cohen et al. 2008; Cohen and van Gaal 2014; Cavanagh et al. 2009; Cavanagh and Frank 2012; Yordanova et al. 2004b; Nigbur et al. 2012; McDermott et al. 2017; Fusco et al. 2018; Töllner et al. 2017) the medial frontal region appears disengaged from the regulation of such processes during errors in older people (Falkenstein et al. 2001). Importantly, however, the increased theta TOTP in the contralateral hemisphere analyzed here was associated with speeded reactions for both correct and incorrect responses of older subjects, similar to young adults who manifest the same relationships at the medial frontal region (Yordanova et al. 2020). Hence, it can be suggested that the functionality of a distributed theta network involved in action monitoring and control is preserved with aging. The lateralized error-related enhancement of theta TOTP may therefore represent an aging-related functional re-distribution of the theta network, with specific inhibition of the key medial frontal mechanisms during error processing.

Another major observation was that the spatial and temporal synchronizations of motor theta oscillations at pre-motor/motor areas contra-lateral to the responding hand were strongly suppressed by errors. It has been demonstrated by cellular and neuroimaging studies that the contra-lateral premotor cortex has a central role in the selection of movements for execution, especially in the selection of learned motor associations (Kalaska and Crammond 1995; Schluter et al. 1998, 2001; Thoenissen et al. 2002; rev. Rushworth et al. 2003). In this regard, the currently found suppression of pre-motor theta synchronization to errors infers the contribution of movement selection processes to error generation in older individuals. Notably, in young adults, pre-motor theta synchronization also was modulated by errors, being however substantially enhanced (Yordanova et al. 2023; Fig. 2B,C - right panel). In line with the established functional relevance of premotor-motor coupling for performance enhancement in both young and old subjects (Michely et al. 2018), the present observations confirm the engagement of pre-motor areas in error generation and further show that the implicated processes of movement selection may play differential roles for incorrect performance in young and old adults.

In older adults, the contra-lateral premotor suppression of temporal and spatial synchronization was accompanied by a similar suppression at the midline fronto-central electrode suggesting a co-reactivity of the two areas during error generation. However, correlational analyses revealed a linked suppression only for left-hand errors and a lack of correlated reactivity for right-hand errors. Hence, during error generation, the synchronization of pre-motor regions may be associated with a separable mechanism of response selection (Rushworth et al. 2003) different from the evaluative control functions of the medial frontal cortex (Vidal et al. 2022; Fu et al. 2023). Indeed, the types of errors analyzed here were different for the left and the right hand. While the speeded left-hand errors appear associated with disinhibition, the tendency for slower right-hand errors may rather reflect a difficulty in motor response selection and execution. This functional asymmetry of error types may have contributed to the differential co-reactivity of the medial frontal and pre-motor regions.

Next, the present results show that despite the lack of enhanced theta oscillations at the medial frontal region in older people, the medial frontal cortex was synchronized with contra-lateral sensorimotor regions during correct motor response generation. This result points to the preserved maintenance of communications between these areas in older subjects perhaps reflecting preserved continuous feedback from sensorimotor areas and monitoring of ongoing movements (Urbano et al. 1998a, b; Töllner et al. 2017). It cannot be excluded that the suppressed synchronization between the medial frontal and sensorimotor regions in old adults is a reflection of the overall decoupling of the medial frontal region during errors. However, this synchronization was inhibited during errors also in young adults, although the medial frontal connectivity was not suppressed. (Yordanova et al. 2023; Fig. 2D). It can be therefore suggested that the error-related disconnection between sensorimotor and fronto-medial areas reflects a more general mechanism of error processing not specifically linked to aging. Yet, in young adults, the connections were inhibited for only slow right-hand errors and not for fast left-hand errors. (Yordanova et al. 2023; Fig. 1 and 2D - right panel). Despite the overall slowing in older adults, they manifested the same asymmetric performance pattern but connections were inhibited for errors with each hand. Together, the observations from the two age groups suggest that error generation/monitoring controlled by the medial frontal cortex may be different for fast and slow motor reactions. Alternatively, they may reflect an aging-related functional asymmetry in processing left-hand motor actions. This possibility aligns with the higher error rate for the left hand observed here as well as with previously established alterations in motor oscillations of correct left-hand responses of older individuals (Yordanova et al. 2020).

Another main result of the present study was the suppressed connectivity and temporal phase-synchronization of motor theta networks in the posterior left hemisphere of older adults, irrespective of the hand with which the error was made. Existing neurocognitive models propose a core function of the lateral parietal activity for mediating an “integrative hub” that combines bottom-up multimodal inputs with top-down controlling signals (Cabeza et al. 2008, 2012; Humphreys et al. 2017; Seghier 2013). Gottlieb et al. (2009) also have suggested that in the lateral parietal area, attentional “priority maps” are formed, representing the dynamic selection of environmental bottom-up features by the top-down focus of attention. In a recent review, Friedman and Ibos (2018) provide further evidence that by encoding sensory, cognitive, and motor-related signals during a wide range of contexts, the posterior parietal cortex serves as a central interface in which these signals converge in order to sharpen behaviorally relevant stimuli and to adaptively influence specific motor networks. Importantly, in a series of studies, Rushworth et al. have demonstrated the critical role of the left parietal cortex for the control of motor attention. They have shown that damage in the left parietal cortex impairs the ability to shift the focus of motor attention from one movement to the next (Rushworth et al. 1997). The application of Positron Emission Tomography (PET) and repetitive Transcranial Magnetic Stimulation (rTMS) has additionally confirmed that the mechanisms of motor attention are lateralized to the left parietal hemisphere in humans (Rushworth et al. 2001a, b, 2003). In addition, Davranche et al. (2011) have found that temporal orienting (i.e., orienting attention to the moment of event occurrence) selectively activates the left parietal cortex for both motor and visual events. In view of these reports, the error-related suppression of connectivity and temporal synchronization in left posterior areas observed here suggests that in older adults, a deficiency in the integration of top-down with bottom-up signals supporting the dynamic focus of motor attention may contribute to incorrect motor reactions. Such a deficit may affect the control and monitoring of actions by the medial frontal cortex focused at FCz, which is in line with the error-related reduction of the FCz-guided synchronization with left parietal regions for each hand. Since no such error-related effects are detected in the left posterior regions of young adults (Yordanova et al. 2023; Fig. 2B,C,D - right panel), the implicated impairment of dynamic motor attention appears to emerge as a function of increasing age in humans.

The present study has several limitations. First, analyses were designed to guarantee a reliable evaluation of error signals by controlling the number of trials (signal-to-noise ratio) but this approach reduced the sample size. Larger sample sizes are needed to support the current findings. Another limitation was that the temporal evolution of motor theta parameters was not assessed, and accordingly, conclusions about the relationships between possible mechanisms and inter-regional interactions are implicative. As a future direction of research, it would be most relevant to establish which of the suggested correlates of error processing may be regarded as precursors of errors or as signals subserving evaluative post-error processes.

## Supporting information

Results of young adults

## 5. Conclusions

The results of the present study demonstrate that error generation in old individuals is accompanied by a substantial suppression of the temporal and spatial synchronization of motor theta oscillations at extended cortical locations involving the medial frontal and contra-lateral premotor and left posterior regions. These observations demonstrate that a distributed theta network is engaged in error generation and processing, and imply that difficulties in controlling the focus of motor attention and response selection contribute to performance impairment in aged individuals.

## Data Availability

The datasets used and analyzed during the current study are available from the corresponding author on reasonable request.

## Acknowledgments

Work was supported by the National Research Fund of the Ministry of Education and Science, Sofia, Bulgaria (Project DN13-7/2017).

## References

1. Alexander WH, Brown JW (2011) Medial prefrontal cortex as an action-outcome predictor. Nat Neurosci 14:1338–1344. https://doi.org/10.1038/nn.2921

2. Babiloni F, Babiloni C, Carducci F, Fattorini L, Onorati P, Urbano A (1996) Spline Laplacian estimate of EEG potentials over a realistic magnetic resonance-constructed scalp surface model. Electroencephalogr Clin Neurophysiol 98:363–373. https://doi.org/10.1016/0013-4694(96)00284-2

3. Band G, Kok A (2000) Age effects on response monitoring in a mental-rotation task. Biol Psychol 51:201–221. https://doi.org/10.1016/s0301-0511(99)00038-1

4. Botvinick MM, Braver TS, Barch DM, Carter CS, Cohen JD (2001) Conflict monitoring and cognitive control. Psychol Rev 108:624–652. https://doi.org/10.1037/0033-295x.108.3.624

5. Botvinick MM, Cohen JD, Carter CS (2004) Conflict monitoring and anterior cingulate cortex: an update. Trends Cogn Sci 8:539–546. https://doi.org/10.1016/j.tics.2004.10.003

6. Cabeza R (2002) Hemispheric asymmetry reduction in older adults: the HAROLD model. Psychol Aging 17:85–100. https://doi.org/10.1037//0882-7974.17.1.85

7. Cabeza R, Anderson ND, Locantore JK, McIntosh AR (2002) Aging gracefully: compensatory brain activity in high-performing older adults. NeuroImage 17:1394–1402. https://doi.org/10.1006/nimg.2002.1280

8. Cabeza R, Ciaramelli E, Olson IR, Moscovitch M (2008) The parietal cortex and episodic memory: an attentional account. Nat Rev Neurosci 9:613–625. https://doi.org/10.1038/nrn2459

9. Cabeza R, Ciaramelli E, Moscovitch M (2012) Cognitive contributions of the ventral parietal cortex: an integrative theoretical account. Trends Cogn Sci 16:338–352. https://doi.org/10.1016/j.tics.2012.04.008

10. Carter CS, Braver TS, Barch DM, Botvinick MM, Noll D, Cohen JD (1998) Anterior cingulate cortex, error detection, and the online monitoring of performance. Science 280:747–749. https://doi.org/10.1126/science.280.5364.747

11. Cavanagh JF, Frank MJ (2014) Frontal theta as a mechanism for cognitive control. Trends Cogn Sci 18:414–421. https://doi.org/10.1016/j.tics.2014.04.012

12. Cavanagh JF, Cohen MX, Allen JJ (2009) Prelude to and resolution of an error: EEG phase synchrony reveals cognitive control dynamics during action monitoring. J Neurosci 29:98–105. https://doi.org/10.1523/JNEUROSCI.4137-08.2009

13. Cohen MX (2011) Error-related medial frontal theta activity predicts cingulate-related structural connectivity. NeuroImge 55:1373–1383. https://doi.org/10.1016/j.neuroimage.2010.12.072

14. Cohen MX (2014a) A neural microcircuit for cognitive conflict detection and signaling. Trends Neurosci 37:480–490. https://doi.org/10.1016/j.tins.2014.06.004

15. Cohen MX (2014b) Analyzing Neural Time Series Data: Theory and Practice. The MIT Press, Cambridge, Massachusetts.

16. Cohen MX, Donner TH (2013) Midfrontal conflict-related theta-band power reflects neural oscillations that predict behavior. J Neurphysiol 110:2752–2763. https://doi.org/10.1152/jn.00479.2013

17. Cohen MX, Ridderinkhof KR, Haupt S, Elger CE, Fell J (2008) Medial frontal cortex and response conflict: evidence from human intracranial EEG and medial frontal cortex lesion. Brain Res 1238:127–142. https://doi.org/10.1016/j.brainres.2008.07.114

18. Cohen MX, van Gaal S (2014) Subthreshold muscle twitches dissociate oscillatory neural signatures of conflicts from errors. NeuroImage 86:503–513. https://doi.org/10.1016/j.neuroimage.2013.10.033

19. Davranche K, Nazarian B, Vidal F, Coull J (2011) Orienting attention in time activates left intraparietal sulcus for both perceptual and motor task goals. J Cogn Neurosci 23:3318–3330. https://doi.org/10.1162/jocn_a_00030

20. Dehaene S (2018) The error-related negativity, self-monitoring, and consciousness. Perspect Psychol Sci 13:161–165. https://doi.org/10.1177/1745691618754502

21. Duprez J, Gulbinaite R, Cohen MX (2020) Midfrontal theta phase coordinates behaviorally relevant brain computations during cognitive control. NeuroImage 207:116340. https://doi.org/10.1016/j.neuroimage.2019.116340

22. Falkenstein M, Hohnsbein J, Hoormann J, Blanke L (1990) Effects of errors in choice reaction tasks on the ERP under focused and divided attention. In: Brunia CHM, Gaillard AWK, Kok A (eds) Psychophysiological brain research, Tilburg University Press, Tilburg, pp. 192–195

23. Falkenstein M, Hohnsbein J, Hoormann J, Blanke L (1991) Effects of crossmodal divided attention on late ERP components. II. Error processing in choice reaction tasks. Electroencephalogr Clin Neurophysiol 78:447–455. https://doi.org/10.1016/0013-4694(91)90062-9

24. Falkenstein M, Hoormann J, Hohnsbein J (2001) Changes of error-related ERPs with age. Exp Brain Res 138:258–262. https://doi.org/10.1007/s002210100712

25. Freedman DJ, Ibos G (2018) An integrative framework for sensory, motor, and cognitive functions of the posterior parietal cortex. Neuron 97:1219–1234. https://doi.org/10.1016/j.neuron.2018.01.044

26. Fu Z, Sajad A, Errington SP, Schall JD, Rutishauser U (2023) Neurophysiological mechanisms of error monitoring in human and non-human primates. Nat Rev Neurosci 24:153–172. https://doi.org/10.1038/s41583-022-00670-w

27. Fusco G, Scandola M, Feurra M, Pavone EF, Rossi S, Aglioti SM (2018) Midfrontal theta transcranial alternating current stimulation modulates behavioural adjustment after error execution. Eur J Neurosci 48:3159–3170. https://doi.org/10.1111/ejn.14174

28. Gehring WJ, Goss B, Coles MGH, Meyer DE, Donchin E (1993) A neural system for error detection and compensation. Psychol Sci 4:385–390.

29. Gottlieb J, Balan P, Oristaglio J, Suzuki M (2009) Parietal control of attentional guidance: The significance of sensory, motivational and motor factors. Neurobiol Learn Mem 91:121–128. https://doi.org/10.1016/j.nlm.2008.09.013

30. Gratton G, Coles MGH, Donchin E (1983) A new method for off-line removal of ocular artifact. Electroencephalogr Clin Neurophysiol 55:468–484. https://doi.org/10.1016/0013-4694(83)90135-9

31. Hoffmann S, Labrenz F, Themann M, Wascher E, Beste C (2014) Crosslinking EEG time-frequency decomposition and fMRI in error monitoring. Brain Struct Funct 219:595–605. https://doi.org/10.1007/s00429-013-0521-y

32. Holroyd CB, Coles MG (2002) The neural basis of human error processing: reinforcement learning, dopamine, and the error-related negativity. Psychol Rev 109:679–709. https://doi.org/10.1037/0033-295x.109.4.679

33. Humphreys GF, Lambon Ralph MA (2017) Mapping domain-selective and counterpointed domain-general higher cognitive functions in the lateral parietal cortex: Evidence from fMRI comparisons of difficulty-varying semantic versus visuo-spatial tasks, and functional connectivity analyses. Cereb Cortex 27:4199–4212. https://doi.org/10.1093/cercor/bhx107

34. Kalaska JF, Crammond DJ (1995) Deciding not to GO: neuronal correlates of response selection in a GO/NOGO task in primate premotor and parietal cortex. Cereb Cortex 5:410–428. https://doi.org/10.1093/cercor/5.5.410

35. Koen JD, Rugg MD (2019) Neural dedifferentiation in the aging brain. Trends Cogn Sci 23: 547–559. https://doi.org/10.1016/j.tics.2019.04.012

36. Kolev V, Falkenstein M, Yordanova J (2005) Aging and error processing: Time-frequency analysis of error-related potentials. J Psychophysiol 19:289–297. https://doi.org/10.1027/0269-8803.19.4.289

37. Kolev V, Beste C, Falkenstein M, Yordanova J (2009) Error-related oscillations: Effects of aging on neural systems for behavioural monitoring. J Psychophysiol 23:216–223. https://doi.org/10.1027/0269-8803.23.4.216

38. Lachaux JP, Rodriguez E, Martinerie J, Varela FJ (1999) Measuring phase synchrony in brain signals. Hum Brain Mapp 8:194–208. https://doi.org/10.1002/(SICI)1097-0193(1999)8:4%3C194::AID-HBM4%3E3.0.CO;2-C

39. Li SC, Lindenberger U, Sikström S (2001) Aging cognition: from neuromodulation to representation. Trends Cogn Sci 5:479–486. https://doi.org/10.1016/s1364-6613(00)01769-1

40. Liu L, Rosjat N, Popovych S, Wang BA, Yeldesbay A, Toth TI, Viswanathan S, Grefkes CB, Fink GR, Daun S (2017) Age-related changes in oscillatory power affect motor action. PLoS One 12:e0187911. https://doi.org/10.1371/journal.pone.0187911

41. Luu P, Tucker DM (2001) Regulating action: alternating activation of midline frontal and motor cortical networks. Clin Neurophysiol 112:1295–1306. https://doi.org/10.1016/s1388-2457(01)00559-4

42. Mathalon DH, Bennett A, Askari N, Gray EM, Rosenbloom MJ, Ford JM (2003) Response-monitoring dysfunction in aging and Alzheimer’s disease: an event-related potential study. Neurobiol Aging 24: 675–685. https://doi.org/10.1016/s0197-4580(02)00154-9

43. McDermott TJ, Wiesman AI, Proskovec AL, Heinrichs-Graham E, Wilson TW (2017) Spatiotemporal oscillatory dynamics of visual selective attention during a flanker task. NeuroImage 156:277–285. https://doi.org/10.1016/j.neuroimage.2017.05.014

44. Michely J, Volz LJ, Hoffstaedter F, Tittgemeyer M, Eickhoff SB, Fink GR, Grefkes C (2018) Network connectivity of motor control in the ageing brain. NeuroImage Clin 18:443–455. https://doi.org/10.1016/j.nicl.2018.02.001

45. Nieuwenhuis S, Ridderinkhof KR, Talsma D, Coles MG, Holroyd CB, Kok A, van der Molen MW (2002) A computational account of altered error processing in older age: Dopamine and the error-related negativity. Cogn Affect Behav Neurosci 2:19–36. https://doi.org/10.3758/cabn.2.1.19

46. Nigbur R, Cohen MX, Ridderinkhof KR, Stürmer B (2012) Theta dynamics reveal domain-specific control over stimulus and response conflict. J Cogn Neurosci 24: 1264–1274. https://doi.org/10.1162/jocn_a_00128

47. Nunez PL, Srinivasan R, Westdorp AF, Wijesinghe RS, Tucker DM, Silberstein RB, Cadusch PJ (1997) EEG coherency. I. Statistics, reference electrode, volume conduction, Laplacians, cortical imaging, and interpretation at multiple scales. Electroencephalogr Clin Neurophysiol 103:499–515. https://doi.org/10.1016/s0013-4694(97)00066-7

48. Perrin F, Pernier J, Bertrand O, Echallier JF (1989) Spherical splines for scalp potential and current density mapping. Electroencephalogr Clin Neurophysiol 72:184–187. https://doi.org/10.1016/0013-4694(89)90180-6

49. Popovych S, Rosjat N, Toth TI, Wang BA, Liu L, Abdollahi RO, Viswanathan S, Grefkes C, Fink GR, Daun S (2016) Movement-related phase locking in the delta-theta frequency band. NeuroImage 139:439–449. https://doi.org/10.1016/j.neuroimage.2016.06.052

50. Reuter-Lorenz P (2002) New visions of the aging mind and brain. Trends Cogn Sci, 6:394–400. https://doi.org/10.1016/s1364-6613(02)01957-5

51. Rushworth MF, Nixon PD, Renowden S, Wade DT, Passingham RE (1997) The left parietal cortex and motor attention. Neuropsychologia 35:1261–1273. https://doi.org/10.1016/s0028-3932(97)00050-x

52. Rushworth MF, Krams M, Passingham RE (2001a) The attentional role of the left parietal cortex: the distinct lateralization and localization of motor attention in the human brain. J Cogn Neurosci 13:698–710. https://doi.org/10.1162/089892901750363244

53. Rushworth MF, Ellison A, Walsh V. (2001b) Complementary localization and lateralization of orienting and motor attention. Nat Neurosci 4:656–661. https://doi.org/10.1038/88492. Erratum in: Nat Neurosci 2001 4:959.

54. Rushworth MF, Johansen-Berg H, Göbel SM, Devlin JT (2003) The left parietal and premotor cortices: motor attention and selection. NeuroImage 20, Suppl 1, S89–100. https://doi.org/10.1016/j.neuroimage.2003.09.011

55. Schluter ND, Rushworth MF, Passingham RE, Mills KR (1998) Temporary interference in human lateral premotor cortex suggests dominance for the selection of movements. A study using transcranial magnetic stimulation. Brain 121: 785–799. https://doi.org/10.1093/brain/121.5.785

56. Schluter ND, Krams M, Rushworth MF, Passingham RE (2001) Cerebral dominance for action in the human brain: the selection of actions. Neuropsychologia 39: 105–113. https://doi.org/10.1016/s0028-3932(00)00105-6

57. Seghier ML (2013) The angular gyrus: Multiple functions and multiple subdivisions. Neuroscientist 19:43–61. https://doi.org/10.1177/1073858412440596

58. Silvetti M, Seurinck R, Verguts T (2011) Value and prediction error in medial frontal cortex: integrating the single-unit and systems levels of analysis. Front Hum Neurosci 5:75. https://doi.org/10.3389/fnhum.2011.00075

59. Tallon-Baudry C, Bertrand O, Delpuech C, Permier J (1997) Oscillatory gamma-band (30-70 Hz) activity induced by a visual search task in humans. J Neurosci 17:722–734. https://doi.org/10.1523/JNEUROSCI.17-02-00722.1997

60. Thoenissen D, Zilles K, Toni I (2002) Differential involvement of parietal and precentral regions in movement preparation and motor intention. J Neurosci 22:9024–9034. https://doi.org/10.1523/JNEUROSCI.22-20-09024.2002

61. Töllner T, Wang Y, Makeig S, Müller HJ, Jung TP, Gramann K (2017) Two independent frontal midline theta oscillations during conflict detection and adaptation in a Simon-type manual reaching task. J Neurosci 37: 2504–2515. https://doi.org/10.1523/JNEUROSCI.1752-16.2017

62. Ullsperger M, von Cramon DY (2001) Subprocesses of performance monitoring: a dissociation of error processing and response competition revealed by event-related fMRI and ERPs. NeuroImage 14:1387–1401. https://doi.org/10.1006/nimg.2001.093

63. Ullsperger M, von Cramon DY (2004) Neuroimaging of performance monitoring: error detection and beyond. Cortex 40:593–604. https://doi.org/10.1016/s0010-9452(08)70155-2

64. Urbano A, Babiloni C, Onorati P, Babiloni F (1996) Human cortical activity related to unilateral movements. A high resolution EEG study. NeuroReport 8:203–206. https://doi.org/10.1097/00001756-199612200-00041

65. Urbano A, Babiloni C, Onorati P, Carducci F, Ambrosini A, Fattorini L, Babiloni F (1998a) Responses of human primary sensorimotor and supplementary motor areas to internally triggered unilateral and simultaneous bilateral one-digit movements. A high-resolution EEG study. Eur J Neurosci 10:765–770. https://doi.org/10.1046/j.1460-9568.1998.00072.x

66. Urbano A, Babiloni C, Onorati P, Babiloni F (1998b) Dynamic functional coupling of high resolution EEG potentials related to unilateral internally triggered one digit movements. Electroencephalogr Clin Neurophysiol 106:477–487. https://doi.org/10.1016/s0013-4694(97)00150-8

67. Varela F, Lachaux JP, Rodriguez E, Martinerie J (2001) The brainweb: phase synchronization and large-scale integration. Nat Rev Neurosci 2:229–239. https://doi.org/10.1038/35067550

68. Vidal F, Burle B, Hasbroucq T (2022) On the comparison between the Nc/CRN and the Ne/ERN. Front Hum Neurosci 15:788167. https://doi.org/10.3389/fnhum.2021.788167

69. Yordanova J, Falkenstein M, Hohnsbein J, Kolev V (2004) Parallel systems of error processing in the brain. NeuroImage 22:590–602. https://doi.org/10.1016/j.neuroimage.2004.01.040

70. Yordanova J, Kolev V, Verleger R, Heide W, Grumbt M, Schürmann M (2017) Synchronization of fronto-parietal beta and theta networks as a signature of visual awareness in neglect. NeuroImage 146:341–354. https://doi.org/10.1016/j.neuroimage.2016.11.013

71. Yordanova J, Falkenstein M, Kolev V (2020) Aging-related changes in motor response-related theta activity. Int J Psychophysiol 153:95–106. https://doi.org/10.1016/j.ijpsycho.2020.03.005

72. Yordanova J, Falkenstein M, Kolev V (2023) Motor oscillations reveal new correlates of error processing in the human brain. Research Square. https://doi.org/10.21203/rs.3.rs-3030180/v1

