## Supplementary material for "A distributed theta network of error generation and processing in aging": Results of young adults

**Supplementary Information**

**Results of young adults (details in Yordanova et al. 2023)**

**Performance**

There was, however, a highly significant Accuracy x Side interaction (F(1/9) = 69.7, p < 0.001). Consistent with the right-handedness of the subjects, RTs were faster for correct right- than left-hand responses (mean 453 ms vs. 471 ms; Side, F(1/9) = 8.9, p = 0.01). For the faster right hand, reactions were slower for error than correct responses (468 vs. 453 ms, p = 0.04), whereas, for the slower left hand, reactions were faster for error than correct responses (460 vs. 471 ms, p = 0.05) – Fig. 1 right panel.

**Theta**

(A) Theta TOTP (Figure 2 - right panel).

Theta TOTP was focused on the midline fronto-central region for both the correct and error responses (Region, F(2/18) = 19.6/32.2, p < 0.001; Laterality, F(2/18) = 6.2/6.9, p = 0.01; Region x Laterality, F(4/36) = 9.4/6.3, p = 0.001/0.01 for left-/right-hand responses, respectively).

For errors, there was a substantial TOTP increase (Accuracy, F(1/9) = 56.3/15.4, p < 0.0001/0.004), which was, however, exclusively pronounced at the midline fronto-central electrode FCz (Region x Accuracy F(2/18) = 12.3/12.1, p = 0.001; Laterality x Accuracy, F(2/18) = 3.8/13.7, p = 0.05/<0.001; Region x Laterality x Accuracy, F(4/36) = 7.4/3.4, p = 0.003/0.04). Testing simple Accuracy effects at single electrodes confirmed the error-related power increase at all frontal-central and central locations (F(1/9) = 5.1 – 20.1, p = 0.05 – 0.001), which was most expressed at FCz (F(1/9) = 36.8, p < 0.0001).

(B) Theta PLF (Figure 2 - right panel).

Theta PLF was significantly larger at contra- than ipsilateral regions (Laterality, F(2/18) = 29.8/26.5, p < 0.0001 for left-and right-hand responses, respectively).

Errors were associated with an increase of PLF at pre-motor regions accompanied by a decrease at motor and sensorimotor regions (Accuracy x Region, F(2/18) = 4.01/3.9, p = 0.05). As dynamic maps imply, pre-motor PLF enhancement involved predominantly the ipsi-lateral hemisphere, whereas the sensorimotor PLF reduction involved critically the contra-lateral hemisphere. This error-related topographic re-distribution did not induce significant effects at single electrodes (F(1/9) < 4.9, p > 0.05).

(C) Theta R-PLV (Figure 2 - right panel).

During both correct and error responses, region-specific connectedness with all other cortical areas was significantly stronger for the hemisphere contralateral to the response (Laterality, F(2/18) = 36.3/22.9, p < 0.001), with pre-motor regions being most strongly connected with other cortical regions in the theta band (Region, F(2/18) = 7.1/3.9, p = 0.02/0.05).

Theta R-PLV did not depend significantly on whether the generated response was correct or incorrect (main and interactive Accuracy for each response side, p > 0.05). The maximal expression of R-PLV was after the response and occurred earlier after error (mean±SE = 26±9.6 ms) than after correct responses (mean±SE = 49.9±6.7 ms; Accuracy F(1/9) = 5.9/5.4, p = 0.04/0.05).

(D) Theta FCz-PLV (Figure 2 - right panel).

During both correct and error responses, theta activity at FCz electrode was most strongly synchronized with central and centro-parietal regions of the hemisphere contralateral to the response (Region, F(2/18 = 22.1/12.2, p < 0.001; Laterality, F(1/9) = 46.7/28.5, p < 0.001; Region x Laterality, F(2/18) = 5.04/6.9, p = 0.03/0.006).

Only for the right hand were errors associated with a reduction of FCz-guided theta synchronization at motor and sensorimotor regions of the contra-lateral left hemisphere (Accuracy, F(1/9) = 4.9, p = 0.05; Accuracy x Region, F(2/18) = 3.9, p = 0.05; Accuracy x Laterality, F(1/9) = 8.9, p = 0.01; Accuracy x Region x Laterality, F(2/18) = 14.1, p = 0.001). Accordingly, only for right-hand responses were simple Accuracy effects significant at C3 and CP3 electrodes (F(1/9) = 6.02 – 18.2, p = 0.04 – 0.003).

**References**

1. Yordanova J, Falkenstein M, Kolev V (2023) Motor oscillations reveal new correlates of error processing in the human brain. Research Square. <https://doi.org/10.21203/rs.3.rs-3030180/v1>
